# Adolescent obesity induces sex-specific alterations of action control

**DOI:** 10.64898/2026.01.29.702500

**Authors:** Diptendu Mukherjee, Solenne Rougeux, Robert T. West, Ahlima Roumane, Kate Z. Peters, Fabien Naneix

## Abstract

The prevalence of obesity is rising worldwide in young people and is associated with poor long-term health outcomes. To counter obesity, weight loss strategies especially involve changes in feeding behaviors and food choice. However, the high level of relapse to unhealthy dietary habits represents an important challenge, suggesting long-term alterations of decision-making and food-seeking processes. Previous studies showed that adolescence is critical for the development of decision-making functions. Thus, it is essential to understand the precise impact of the exposure to obesogenic diets during this life stage on the different processes underlying flexible control of food-seeking actions.

To address this, we gave mice access to high-fat diets (HFDs) with different fat contents during adolescence and investigated the long-lasting impact on action control at adulthood after a switch to a healthy diet. We uncovered important sex differences. In both males and females, exposure to HFD with very high-fat content (60%) promoted habitual behavior, which is less flexible to adapt to changes in outcome value or action-outcome relationships. In contrast, exposure to HFD with lower fat content (45%) impaired action control based on the updating of outcome value in males only, while impairing action control based on the updating of action-outcome relationships in females only. These findings highlight how the consumption of obesogenic diets during adolescence has long-lasting, diet- and sex-dependent effects on decision-making processes, promoting habitual responses to food. These changes may support long-term vulnerability for mental and physiological health conditions.

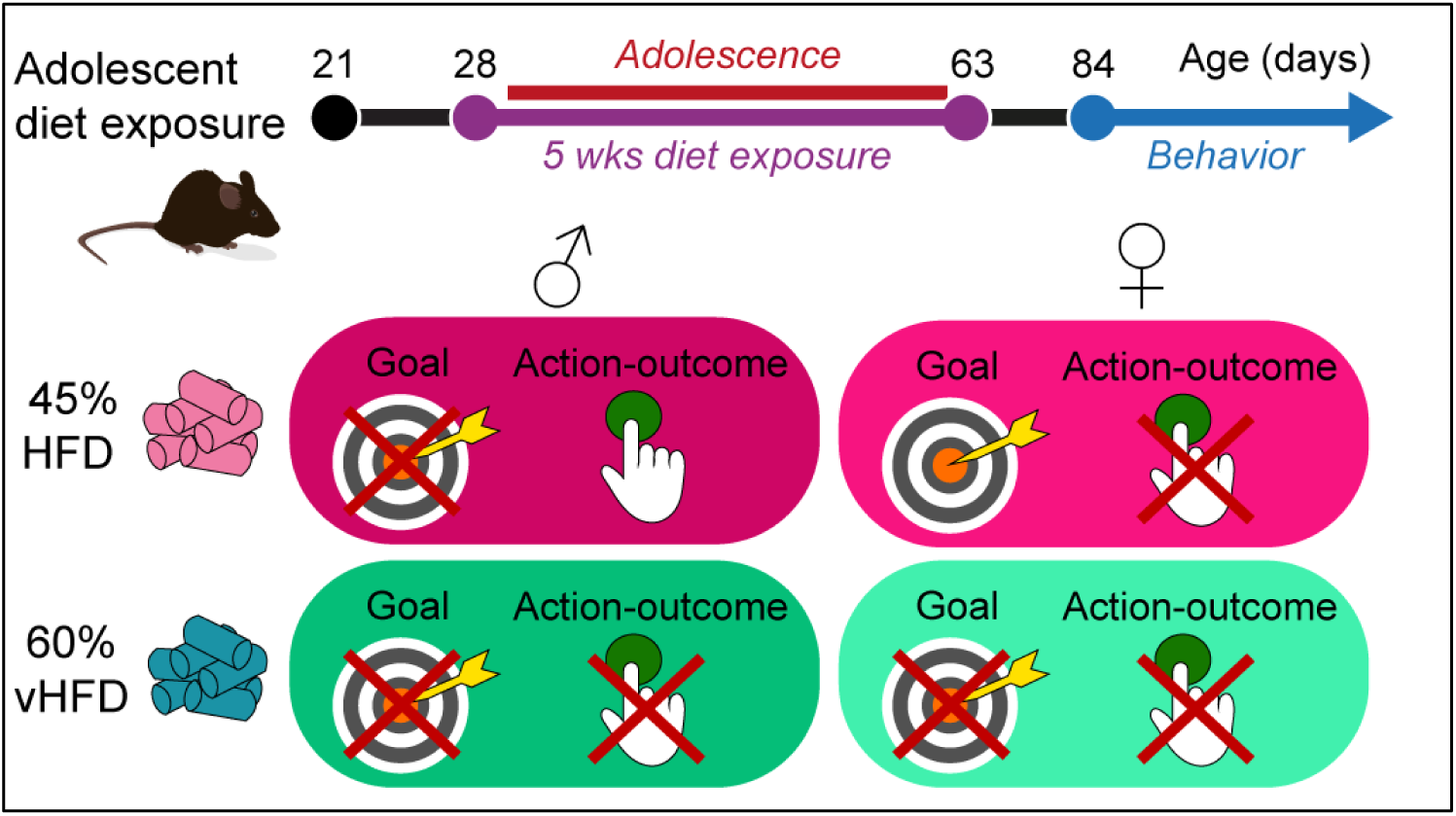

## INTRODUCTION

Obesity and overweight are among the major health challenges for current societies with more than 50% of adults worldwide considered overweight or obese. Recently, the rate of obesity has particularly risen amongst teenagers, quadrupling since 1990, with a similar prevalence between sexes [1]. Obesity in childhood and adolescence is a major predictor of adult obesity. Adolescence is also a critical window for cognitive development, especially for processes controlling feeding and value-based decision-making [2–7]. Thus, it is critical to understand the potential long-term and long-lasting impact of obesogenic diets on these processes.

The overconsumption of highly palatable, energy-dense foods represents a major driver of obesity, leading to metabolic dysfunctions (e.g. cardiovascular disorders, diabetes) as well as neurobiological and cognitive alterations [8]. Most studies focused mainly on diet-induced changes in the brain regulation of energy balance [9,10]. However, weight-loss strategies, such as diet changes and increased exercise, often show poor long-term outcomes on body weight associated with a relapse to unhealthy dietary and lifestyle habits [11,12]. A possibility to explain this may be an alteration of decision-making processes by obesogenic diets [8,13].

Adaptive flexible control of food-seeking requires learning the relationships between actions and their consequences, while constantly monitoring the value of these outcomes [14–19]. In contrast, habitual responses arise when behavior becomes less flexible and not goal-directed anymore, occurring independently of the action consequences or of the outcome value [15,16,18]. An imbalance toward this habitual system may lead to automatic responding for food, contributing to overconsumption and poor food choice. Several studies in humans and preclinical models show that consumption of highly palatable diets at adulthood is associated with reduced behavioral flexibility and habitual control of food-seeking [20–25]. However, most of these studies focused on control based on outcome value, overlooking other cognitive processes such as the monitoring of changes in action-outcome (A-O) causal relationships essential to flexibly adapt and explore alternative behavioral strategies [18,19]. Moreover, we and others have shown that a similar diet can have greater impact if exposure occurs during key developmental windows like adolescence [26–29]. Given the rising rates of obesity in teens, understanding the impact of obesogenic diets during adolescence on the cognitive processes controlling food choice and food-seeking is essential to develop better therapeutic approaches.

Increasing evidence points at sex differences in multiple cognitive processes including decision-making [30–33]. As decision-making processes are still under development during early life [4–7,34–36] and show key puberty-related changes [5,36], it is crucial to understand the long-lasting impact of adolescent obesogenic diets on action control processes [37,38].

Here, we investigated the long-lasting impact of exposure to obesogenic diets during adolescence on goal-directed and habitual control of actions later in life. In contrast to most studies we limited exposure to adolescence to isolate long-term effects on cognition and behavior independently of ongoing neuronal and metabolic alterations associated with chronic overweight and obesity. To compare directly the effects of different levels of diet-induced obesity on cognitive functions, we used two high-fat diets known to induce different levels of weight gain [39,40]. We show that exposure to diets with higher fat content alters goal-directed behavior based on either outcome’s value or action-outcome (A-O) causal relationships in both males and females. In contrast, diets with “moderate” fat content selectively alter action control based on outcome value in males, while selectively altering action control based on A-O relationships in females. These findings demonstrate for the first time important sex and nutrient specific effects of adolescent exposure to obesogenic diet on value-based decision-making which endure into adulthood.

## MATERIAL AND METHODS

### Subjects

Male and female C57Bl/6J mice were bred at the Medical Research Unit (University of Aberdeen). At weaning (postnatal day, PND, 21) mice were housed in polycarbonate stock cages (450 x 280 x 130 mm, Techniplast 1292N; 4-7 mice per cage according to their sex) for the duration of diet exposure. For behavioral testing, they were housed in polycarbonate cages (369 x 165 x 132 mm, Techniplast 1145; 1-4 mice per cage). Environmental enrichment was provided by tinted polycarbonate tubes and nesting material. They were maintained in a temperature (19-22°C) and humidity (45-55%) controlled environment with a 12 h light/dark cycle (lights on at 7:00 AM) and with food and water available *ad libitum*, except when otherwise stated. All behavioral experiments took place during the light phase. All procedures were performed in accordance with the Animals (Scientific Procedures) Act 1986 and carried out under Project License P58595877.

### Diets

Mice were randomly assigned to one of the three diet conditions for which they were exposed for 5 weeks between PND28 and PND63 (**Figure 1A**) [2,41]: **1)** Standard diet (SD group) with only access to standard chow diet (3.6 kcal/g; 69% kcal from carbohydrate, 9% lipids, 22% proteins; CMR, SDS), **2)** High-fat diet (HFD group) with combined access to standard chow and to 45% fat diet (4.7 kcal/g; 35% kcal from carbohydrate, 45% lipids mainly from lard, 20% proteins; D12451, Research Diets), **3)** Very High-Fat diet (vHFD group) with combined access to standard chow and to 60% fat diet (5.2 kcal/g; 20% kcal from carbohydrate, 60% lipids mainly from lard, 20% proteins; D12492, Research Diets; **Supplemental Table 1**). All diets were provided *ad libitum*. Animals’ body weight, water and food intake were recorded weekly. At PND63, all mice were switched to standard chow diet and water only. Food restriction started two weeks after the end of diet exposure (> PND77). Mice were progressively food restricted to ∼85-90% of their free-feeding body weight over a week before the start of behavioral testing. HFD and vHFD body weights were compared to the average initial body weight of their appropriate control groups. Mice were always weighed and fed once all testing of the day was completed. The amount of food was adjusted based on individual body weight and the amount of food earned each day (both in terms of g and kcal).

**Figure 1.**
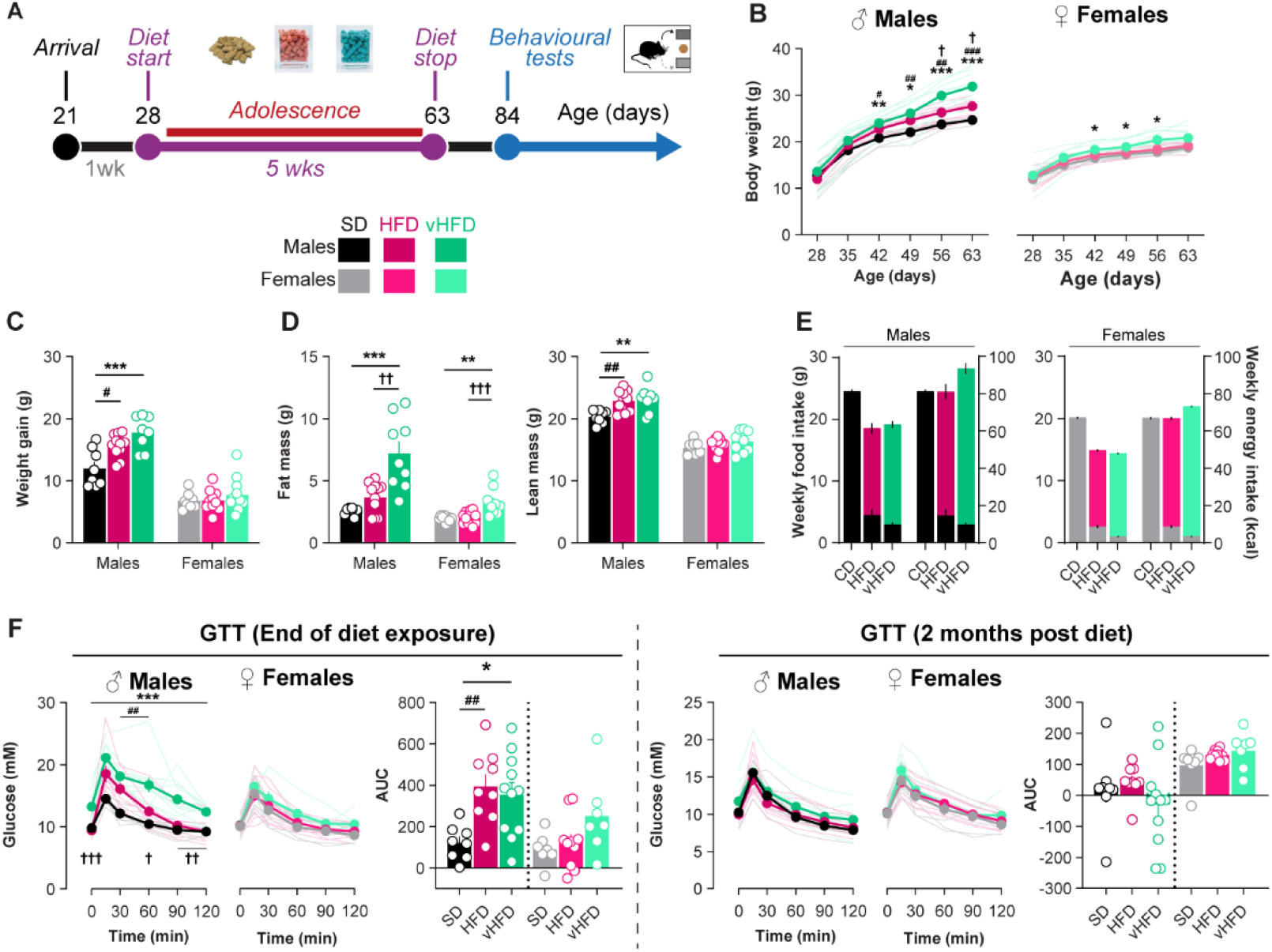
Obesogenic diets during adolescence induce sex-specific impact on body weight and metabolic health. **A.** Schematic representation of experimental design. Mice had continuous access to either standard chow diet only (SD, 9% kcal from fat) or simultaneous access to SD and either high-fat diet (HFD, 45% kcal from fat) or very high-fat diet (vHFD, 60% kcal from fat) during adolescence (postnatal days 28–63). At adulthood, all mice had only access to SD for two weeks before being mildly food restricted and the start of behavioral testing. **B-D.** HFD and vHFD exposure during adolescence differentially impacted body weight (**B**; two-way ANOVA; Males: Age F_(1.87, 46.79)_ = 608.4, p < 0.001, η_p_^2^ = 0.96; Diet F_(2,25)_ = 8.5, p = 0.001, η_p_^2^ = 0.41; Age x Diet F_(3.74, 46.79)_ = 9.6, p < 0.001, η_p_^2^ = 0.43; Bonferroni’s *post hoc* tests SD vs HFD p < 0.05 from P56, SD vs vHFD p < 0.05 from P42, HFD vs vHFD p < 0.05 from P56 / Females: Age F_(1.75, 45.39)_ = 277.2, p < 0.001, η_p_^2^ = 0.91; Diet F_(2,26)_ = 4.9, p = 0.01, η_p_^2^ = 0.27; Age x Diet F_(3.49, 45.39)_ = 1.6, p = 0.2 η_p_^2^ = 0.11; Bonferroni’s *post hoc* tests SD vs HFD all p > 0.1, SD vs vHFD p < 0.05 from P42, HFD vs vHFD all p > 0.06), weight gain (**C**; one-way ANOVA; Males: F_(2,25)_ = 12.4, p < 0.001, η^2^ = 0.5; Bonferroni’s *post hoc* tests SD vs HFD p = 0.02, SD vs vHFD p < 0.001, HFD vs vHFD p = 0.1 / Females F_(2,26)_ = 1.2, p = 0.3, η^2^ = 0.09) and body composition (fat and lean mass, **D**; one-way ANOVA; Males: Fat mass F_(2,25)_ = 15.8, p < 0.001, η^2^ = 0.56, Bonferroni’s *post hoc* tests SD vs HFD p = 0.6, SD vs vHFD p < 0.001, HFD vs vHFD p < 0.001; Lean mass F_(2,25)_ = 7.5, p = 0.003, η^2^ = 0.38, Bonferroni’s *post hoc* tests SD vs HFD p = 0.009, SD vs vHFD p = 0.005, HFD vs vHFD p = 1.0 / Females: Fat mass F_(2,26)_ = 10.7, p < 0.001, η^2^ = 0.50, Bonferroni *post hoc* tests SD vs HFD p = 1.0, SD vs vHFD p = 0.002, HFD vs vHFD p < 0.001; Lean mass F_(2,26)_ = 1.3, p = 0.3, η^2^ = 0.09) in male and female mice. **E.** HFD and vHFD groups showed lower amounts of food intake (g) but slightly elevated energy intake (kcal, estimated from cage consumption), with a clear preference for obesogenic diets (pink/green) compared to standard diet (black/grey). **F.** HFD- and vHFD male, but not female, mice showed an impaired glucose clearance at the end of diet exposure (*left*; one-way ANOVA; Males: AUC F_(2, 25)_ = 6.6, p = 0.005, η^2^ = 0.34; Bonferroni’s *post hoc* tests SD vs HFD p = 0.008, SD vs vHFD p = 0.02, HFD vs vHFD p = 1.0 / Females: AUC F_(2, 21)_ = 2.7, p = 0.09, η^2^ = 0.21) but not 2 months after (*right*; one-way ANOVA; Males: AUC F_(2, 25)_ = 1.1, p = 0.3, η^2^ = 0.08 / Females: AUC F_(2, 21)_ = 2.0, p = 0.2, η^2^ = 0.16). SD, 8M/7-8F (black/gray respectively); HFD, 9-11M/10-12F (dark and light pink); vHFD, 9-11M/7-9F (dark and light green). Data are presented as mean ± SEM (bars or closed circles) with individual values (open circles or thin lines). *, **, *** p < 0.05, 0.01 and 0.001 respectively SD vs vHFD. #, ##, ### p < 0.05, 0.01, and 0.001 respectively SD vs HFD; †, ††, ††† p < 0.05, 0.01 and 0.001 respectively HFD vs vHFD (one-way or two-way ANOVA followed by Bonferroni’s *post hoc* tests). Full statistical reporting is provided in **Supplemental Table 2**. Mouse illustration from NIAID NIH BioArt Source (bioart.niaid.nih.gov/bioart/20).

### Body composition analysis

At PND63, the impact of different diets on body composition (fat and lean mass) was determined using EchoMRI^TM^. Mice were weighed before and then gently restrained in plastic holders without anesthesia for the duration of the procedure. Measurements were performed in triplicates.

### Glucose tolerance test (GTT)

At PND63 and PND120, glucose clearance was tested using a GTT. Mice were placed in clean cages and food withheld for five hours. Basal glucose levels were determined from tail puncture (AlphaTrak® III, Zoetis). All mice received an injection of D-glucose (2 mg/g, i.p.; Sigma-Aldrich) and blood glucose levels measured at t+15, 30, 60, 90 and 120 minutes. Mice had *ad libitum* access to water throughout the experiment.

### Behavioral apparatus

Animals were tested in individual standard operant chambers (25 × 32 × 25.5 cm, Med Associates), each located inside sound- and light-attenuating wooden chambers (63 × 44 × 40 cm) with inbuilt ventilation fans producing background noise. Each conditioning chamber was equipped with a house light located on the left wall, a pellet magazine located in the center of the right wall, two retractable levers on each side of the magazine and two LEDs above levers. The magazine was equipped with an infrared light to detect head entries and was connected to a pellet dispenser delivering 20 mg food pellets (Rodent Purified pellets, F0071; Bio-Serv). The house light and fan were turned on at the beginning of each daily session and turned off at the end. Conditioning chambers were controlled via a computer running Med-PC IV Software Suite (Med Associates). All behavioral tests were conducted during the light phase of the light/dark cycle (usually between 9am and 3pm, 7 days a week).

#### Behavioral procedures

##### Instrumental training

Instrumental training procedure was adapted from previous studies conducted in rats [29,34,42]. Initially, mice were trained in two sessions to collect food pellets in the magazine (day 1-2; 30-min session). Food rewards were delivered at a random time 60 s schedule. Mice then received 9 daily sessions of instrumental training, during which they learned initial A-O associations (30 rewards or 40 min maximum). During these sessions, the two levers were presented on each side of the magazine. Mice were trained to press one lever (active lever) to obtain the food pellet. Reinforced lever presses were followed by 5 s time-out period during which a cue light located above the active lever and the magazine light were turned on and lever presses did not result in additional reward delivery. Responses on the other lever (inactive lever) had no consequence. The position of the active and inactive lever was counterbalanced between animals. Mice were trained in daily sessions in a fixed ratio 1 (FR1, *i.e.* each response was rewarded) for 3 days before being shifted to a variable interval 15 sec (VI15, *i.e.* outcomes were delivered for the first active lever press made after an average interval of 15 sec) for 3 days, VI30 for 3 days and VI60 for 17 days.

##### Instrumental outcome devaluation tests

Before the outcome devaluation tests, mice were habituated to the individual consumption polycarbonate cages (369 x 165 x 132 mm, Techniplast 1145) located in a different room for 60 min the day before the first test. Mice received 3 outcome devaluation tests by sensory specific satiety. Tests were given after 3 days of VI60 (early), 9 days of VI60 (mid) and 17 days of VI60 (late) based on previous work [43,44]. Each test was organized as follows: mice were given free access to either the food pellet (Devalued condition) or standard chow (Non devalued condition; 4 g placed in a plastic dish) in individual consumption cages for 60 min. Immediately after, mice were tested in a 15 min choice extinction test (unrewarded) in their conditioning chambers. Then, they were placed back in consumption cages for a 15 min consumption test during which they had concurrent access to 2 g of each food (pellet and chow) to confirm the effectiveness and the specificity of the devaluation procedure. After one day of VI60 retraining, the other food was devalued using a similar procedure. The order of devalued food was counterbalanced between mice.

##### Reversal training and test

Following the last outcome devaluation test, mice received 8 days of VI60 training identical to the ones previously described except that the originally active lever became inactive, and the originally inactive lever became active. Following this training, a last outcome devaluation test was conducted, using an identical procedure as previously described.

##### Instrumental contingency degradation

Following the outcome devaluation tests, mice were retrained on VI60 for two days before the start of the contingency degradation procedure. Each diet condition was divided into two groups for the following 7 days. For the first group, the A-O contingency was maintained on the VI60 schedule as during training (Intact contingency). For the other group, the pellets were now randomly delivered throughout the session based on a VI60 schedule but without being linked to the instrumental response (Degraded contingency). Twenty-four hours after the last contingency degradation session, mice were tested in a 15 min choice extinction test in their conditioning chambers.

##### Progressive ratio

The following days, mice were retrained on VI60 to reach similar levels of instrumental responses. Their motivation for the food reward was then tested under a progressive ratio (PR) schedule, during which the number of lever presses required to earn each reward increased exponentially on successive trials [response ratio = (5 × e^(0.2^ ^×^ ^reward^ ^number)^ − 5)], yielding ratios of 1, 2, 4, 6, 9, 12, 15, 20, 25, 32, 40, 50, 62, 77, 95, 118, etc [45]. The test was stopped after a maximum of 2 hours or if an animal did not earn a pellet within 20 min. Total number of lever presses, number of rewards earned, and breakpoint (final response ratio reached) were recorded. Mice were trained on PR until breakpoints stabilized (± one response schedule for three consecutive days with no increasing or decreasing trend).

#### Experimental design and statistical analysis

For outcome devaluation and contingency degradation tests, analyses were restricted to the first 5 mins to prevent contamination from extinction. Statistical analyses were conducted using GraphPad Prism 9 and SPSS (IBM). Figures were created using GraphPad Prism 9 and Adobe Illustrator. Data are represented as mean ±SEM and individual values. Group sizes were estimated based on previous work with these behavioral tasks and to ensure counterbalancing between action learned and test orders. Data were analyzed by one-, two- or three-way analysis of variance (ANOVA) with or without repeated measures. Greenhouse– Geisser correction was applied to mitigate the influence of unequal variance between conditions for repeated measures when necessary. Bonferroni *post hoc* tests corrected for multiple comparisons were performed when appropriate. To specifically investigate response sensitivity to outcome devaluation or contingency degradation, data from the different tests were compared using multiple pairwise comparisons with Bonferroni’s correction restricted to Devaluation or Degradation effects for each diet as *a priori* comparisons. Tests were also analysed using two-way mixed ANOVAs (Diet x Condition) but never revealed significant Diet x Devaluation or Diet x Contingency interactions. All data were initially compared with Sex as factor. Due to initial differences in weight gain and energy intake during the diet exposure, separate analyses for each sex were also systematically performed as planned comparisons. Homogeneity of variance and normality were measured with Levene’s test and Shapiro–Wilk test, respectively. Based on the robustness of ANOVA to slight non-normality [46], we opted to use parametric tests for consistency between experiments. Measures of effect size (eta squared η^2^ or partial eta squared η_p_^2^ for ANOVA with small = 0.01, medium = 0.06 and large = 0.14, Cohen’s d for between-subject contrasts with small = 0.2, medium = 0.5 and large = 0.8) are stated for each comparison. The alpha risk for the rejection of null hypothesis was set at 0.05. Full statistical reporting is provided in **Supplemental Table 2** and **Supplemental Table 3**.

## RESULTS

### Obesogenic diets during adolescence induce sex-specific impact on body weight and metabolic health

We first measured the impact of exposure to different HFDs during adolescence on body weight and body composition (SD: n = 8 males / 8 females; HFD: n = 11/12; vHFD; n = 9/9). Exposure to either HFD or vHFD during adolescence significantly increased male body weight and total weight gain compared to SD group (**Figure 1B-C**). In females, only consumption of vHFD significantly increased body weight (**Figure 1B-C**). Analysis of body composition at the end of the diet exposure phase confirmed a significant increase in fat mass for both male and female vHFD mice, as well as an increased levels of lean mass in HFD and vHFD male mice (**Figure 1D**).

As mice were group-housed, we do not have individual food or water intake data, precluding statistical analysis. Visual inspection showed lower average food intake in HFD and vHFD groups. However, this lower food intake still resulted in similar or even higher energy consumption in HFD and vHFD groups respectively, compared to SD groups (**Figure 1E**). This is consistent with the higher caloric content of the diet and the higher weight gain and fat mass in these animals (see also **Supplemental Figure 1B**). Water intake seemed identical among all groups (**Supplemental Figure 1A**).

Both HFD and vHFD-exposed males showed an impaired glucose metabolism and clearance after 5 weeks of diet, with vHFD group also exhibiting higher glucose levels at baseline. Alteration of glucose clearance is only observed in vHFD-exposed females, consistent with their obesity phenotype, without basal hyperglycemia. Body weight and glucose metabolism alterations were not maintained after a switch to standard diet (**Figure 1F** and **Supplemental Figure 1C**).

### Long-lasting impact of adolescent obesogenic diets on the control of actions by outcome value

At adulthood (>PND70), mice were trained to perform an action (A; press on the active lever) to obtain a specific outcome (O; food pellet). Another lever with no associated outcome was present to control for learning and general activity. Mice received extensive instrumental training using VI reinforcement schedules, previously described to favor the development of habitual responses **(Figure 2A)** [15]. All mice successfully learned the task, progressively increasing their response rate on the active lever (**Figure 2B**), while they decreased their response on the inactive lever (*data not shown*). Food port entries and latencies to collect food were similar between groups (**Supplemental Figure 2A-B**). These results demonstrate that exposure to obesogenic diets during adolescence did not impair initial instrumental learning and discrimination between different actions. Additional progressive ratio testing performed at the end of all behavioral testing also showed similar instrumental performance and breakpoint between groups confirming that adolescent diet exposure did not impair the reinforcing properties of the food rewards used in the task (**Supplemental Figure 2C**).

**Figure 2.**
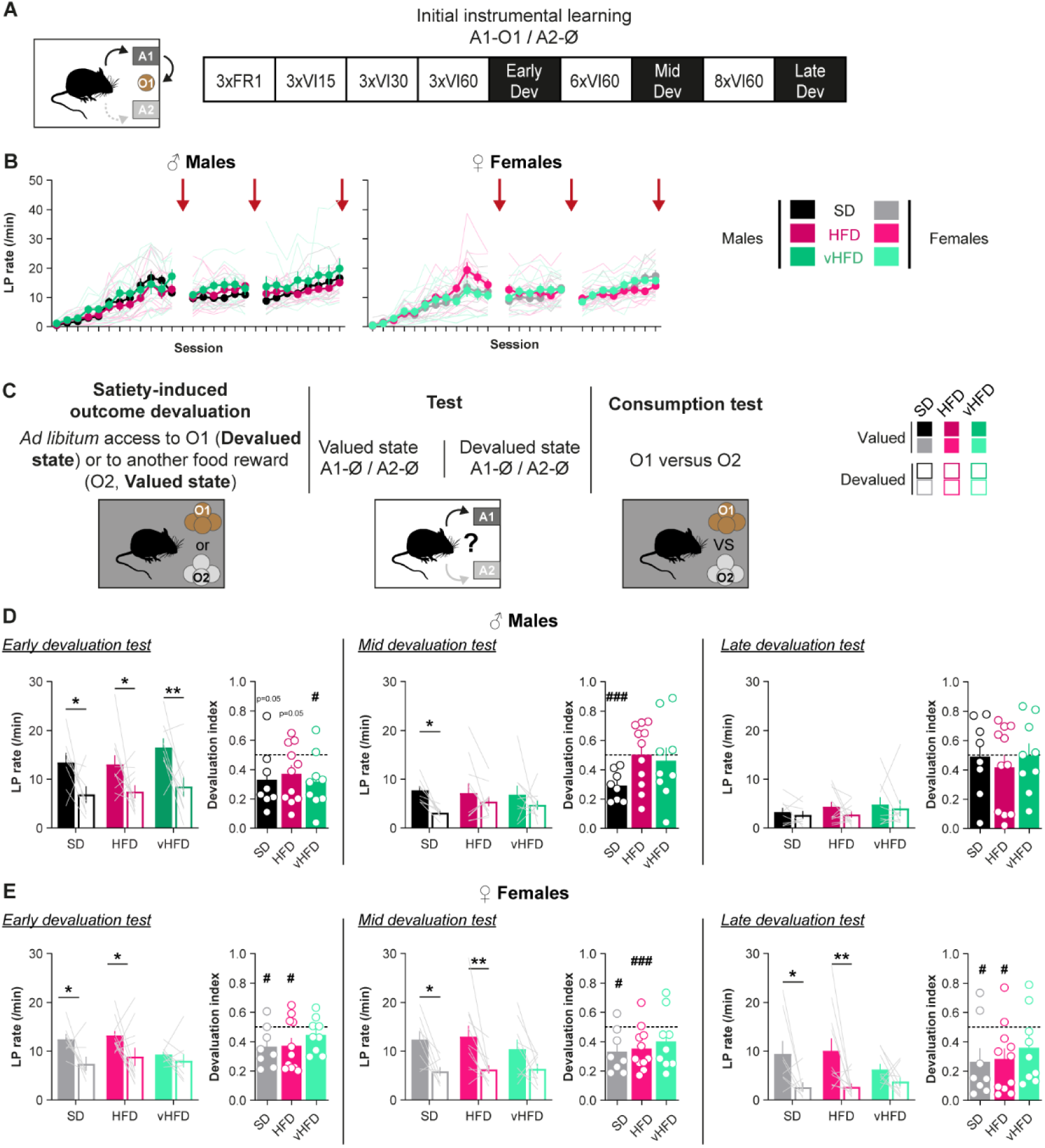
Long-lasting impact of adolescent obesogenic diets on the control of actions by outcome value in a diet- and sex-dependent manner. **A.** Experimental timeline for initial instrumental training and outcome devaluation tests (early, mid and late dev). **B.** Active lever response rate across initial training. Red arrows indicate the timing of the outcome devaluation tests (Males: Diet all Fs < 1.2, all p > 0.3; Session all Fs > 3.5, all p < 0. 001; Diet x Session all Fs < 1.3, p > 0.3 / Females: Diet all Fs < 1.0, all p > 0.4; Session all Fs > 3.7, all p < 0.001; Diet x Session all Fs < 2.6, p > 0.04). **C.** Satiety-induced outcome devaluation procedure. **D-E.** Devaluation test response rate and devaluation index ([Devalued condition response]/[Valued + Devalued conditions responses]) at early (*left*), mid (*middle*) and late (*right*) stages of training for male (**D**; Planned comparisons with Bonferroni’s correction; Early: SD F(1,25) = 4.5, p = 0.04, η_p_^2^ = 0.15; HFD F(1,25) = 4.6, p = 0.04, η_p_ ^2^ = 0.16; vHFD F(1,25) = 7.8, p = 0.01, η_p_ ^2^ = 0.24 / Mid: SD F(1,25) = 5.0, p = 0.03, η_p_ ^2^ = 0.17; HFD F(1,25) = 0.9, p = 0.3, η_p_^2^ = 0.04; vHFD F(1,25) = 1.1, p = 0.3, η_p_^2^ = 0.04 / Late: SD F(1,25) = 0.1, p = 0.7, η_p_^2^ = 0.004; HFD F(1,25) = 1.4, p = 0.3, η_p_^2^ = 0.05; vHFD F(1,25) = 0.2, p = 0.6, η_p_^2^ = 0.009) and female (**E**; Planned comparisons with Bonferroni’s correction; Early: SD F(1,25) = 5.1, p = 0.03, η ^2^ = 0.17; HFD F(1,25) = 5.2, p = 0.03, η_p_^2^ = 0.17; vHFD F(1,25)= 0.4, p = 0.6, η_p_^2^ = 0.01 / Mid: SD F(1,25) = 8.0, p = 0.009, η_p_^2^ = 0.24; HFD F = 11.8, p = 0.002; η_p_^2^ = 0.32; vHFD F(1,25)= 3.3, p = 0.08, η_p_^2^ = 0.12) / Late: SD F(1,25) = 5.5, p = 0.03, η_p_^2^ = 0.18; HFD F(1,25) = 8.9, p = 0.006; η_p_^2^ = 0.26; vHFD F(1,25) = 0.8, p = 0.4, η_p_^2^ = 0.03) mice. SD, 8M/8F (black/gray respectively); HFD, 11M/11F (dark and light pink); vHFD, 9M/9F (dark and light green). Data are presented as mean ± SEM (bars or closed circles) with individual values (thin lines). *, ** p < 0.05 and 0.01 respectively (Planned comparisons Valued vs Devalued with Bonferroni’s correction). #, ### p < 0.05 and 0.001 respectively (one sample t-test versus 0.5). Full statistical reporting is provided in **Supplemental Table 2**. Mouse illustration from NIAID NIH BioArt Source (bioart.niaid.nih.gov/bioart/20).

At three stages of the training their ability to use the current value of the outcome to control their food-seeking behavior was tested, using outcome devaluation by sensory-specific satiety (**Figure 2C**). In all groups, instrumental performance decreased with repeated outcome devaluation tests, despite similar levels between training blocks. In males, the control of instrumental actions progressively shifts from goal-directed (*i.e* sensitive to changes in outcome value) to more habitual (insensitive to outcome devaluation) in the SD group. However, this behavioral shift is observed earlier (after moderate level of training) in both HFD- and vHFD-exposed males (**Figure 2D**).

A different pattern is observed in females. In contrast to the control male SD group, SD females seem to maintain a form of goal-directed control of their actions even after extended training in our experimental conditions (**Figure 2E**). A similar pattern is observed in the HFD-exposed female group. Interestingly, vHFD-exposed females appeared already insensitive to outcome devaluation at early stages of training and maintained this behavioral pattern during the three outcome devaluation tests. Importantly, all mice ate similar amounts during the satiety phase and concurrent access consumption tests, successfully discriminating valued (non-pre-fed) and devalued (pre-fed) foods. This demonstrates the efficiency of the outcome devaluation procedures for all groups (**Supplemental Figure 3A-B**).

Taken together, these results show that exposure to a high-fat diet during adolescence accelerates the transition from goal-directed to habitual control of actions, as shown in response to changes in outcome value. Importantly, our results highlight sex- and diet-related differences with an accelerated transition to habit-like behaviors for all vHFD-exposed groups and in HFD-exposed males, but not female mice.

### Long-lasting impact of adolescent obesogenic diets on the control of actions by action-outcome relationships

We next tested if the exposure to different HFDs during adolescence also impacts flexible control of actions based on action-outcome relationships. For this, we first used an instrumental reversal procedure during which the nature of previously learned active and inactive responses was inverted (**Figure 3A**). All mice quickly switched their response toward the new active lever and decreased their response rate on the new inactive lever/former active lever (**Figure 3B**), suggesting that they were all able to flexibly adapt their instrumental behavior and inhibit their former learning in a reversal setup. We then assessed if mice were able to use this new A-O association to correctly guide their food-seeking when the outcome value is changed (**Figure 3C-E**). The acquisition of a new A-O association was able to restore action control based on outcome value in SD and HFD-exposed males (**Figure 3D**). We also observed a similar, but non-significant, behavioral pattern for the response on the former active lever (**Figure 3E**). However, vHFD-exposed males appeared insensitive to changes in outcome value for both instrumental responses suggesting an impairment in their ability to correctly update and use action-outcome associations. In females, SD control mice correctly showed instrumental responses sensitive to outcome devaluation. A similar pattern is observed on the former active lever (**Figure 3D-E**). Interestingly, HFD-exposed females appeared insensitive to outcome devaluation after response reversal while exhibiting devaluation sensitivity on the former active lever, suggesting a deficit in their ability to correctly update previously learned actions. Finally, vHFD-exposed females appeared as insensitive to outcome devaluation for both actions. As for the first devaluation tests, all mice consumed similar amounts during the satiety phase and then rejected the devalued food during the consumption test (**Supplemental Figure 4**).

**Figure 3.**
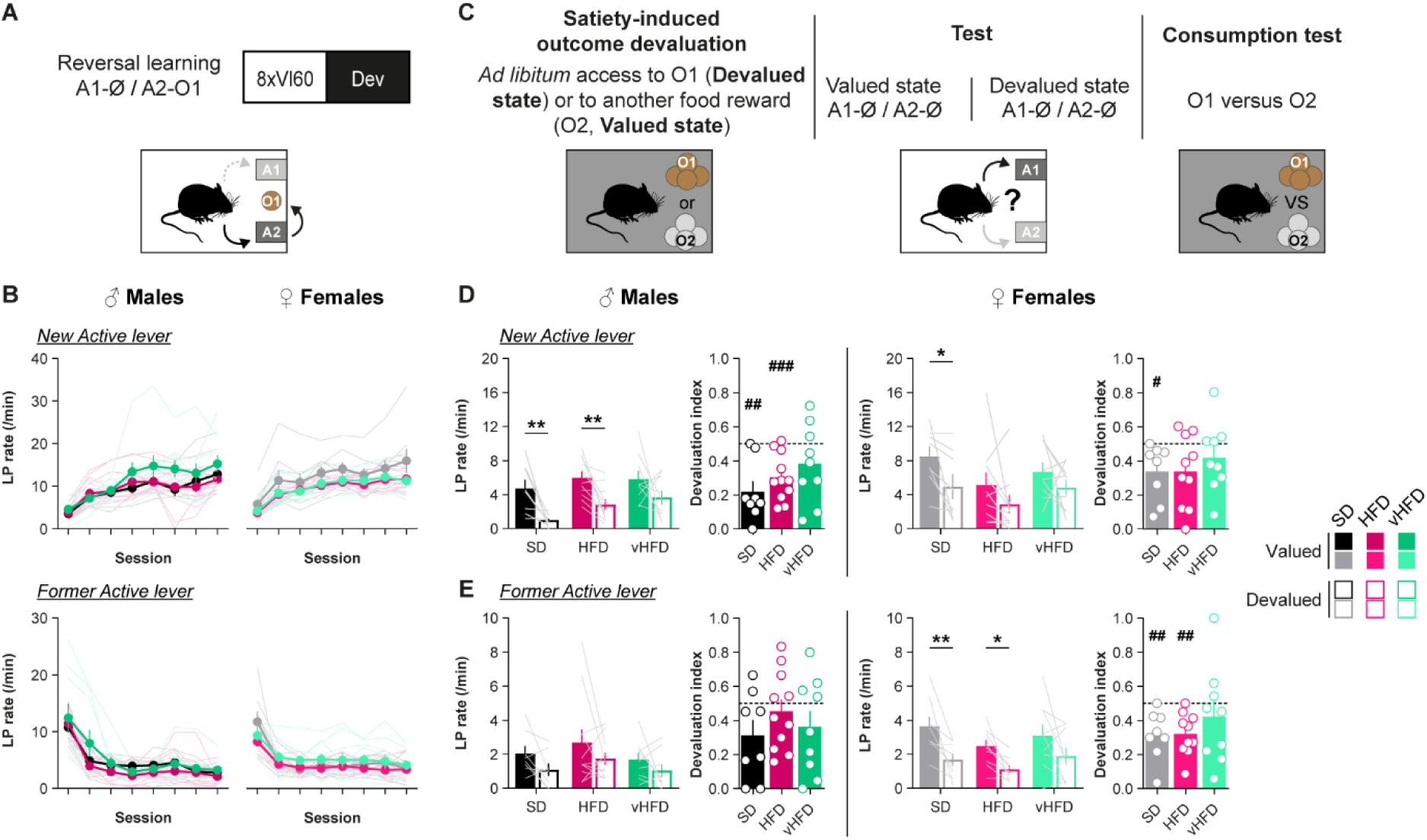
Adolescent obesogenic diets differentially alter action control in response to contingency reversal in a diet- and sex-dependent manner. **A.** Experimental timeline for reversal training and outcome devaluation test. **B.** Response rate on the new active lever (*top*) and the former active lever (*bottom*) for male and female mice (*left and right respectively*; Males: Diet all Fs < 1.4, all p > 0.2; Session all Fs > 30.5, all p < 0.001; Diet x Session all Fs > 1.5, p > 0.2 / Females: Diet all Fs < 2.0, all p > 0.2; Session all Fs > 33.0, all p < 0.001; Diet x Session all Fs < 1.4, p > 0.3). **C.** Satiety-induced outcome devaluation procedure. **D-E.** Devaluation test response rate and devaluation index on the new (**D**; Planned comparisons with Bonferroni’s correction; Males: SD F(1,25) = 8.4, p = 0.008, ηp^2^ = 0.25; HFD F(1,25) = 8.3, p = 0.008, ηp^2^ = 0.25; vHFD F(1,25) = 3.2, p = 0.09, ηp^2^ = 0.11 / Females: SD F(1,24) = 5.0, p = 0.03, ηp^2^ = 0.18; HFD F(1,24)= 2.6, p = 0.1, ηp^2^ = 0.10; vHFD F(1,24)= 1.5, p = 0.2, ηp^2^ = 0.06) and former active lever (**E**; Planned comparisons with Bonferroni’s correction; Males SD F(1,25) = 1.8, p = 0.2, η_p_ ^2^ = 0.07; HFD F(1,25) = 2.4, p = 0.1, ηp^2^ = 0.09; vHFD F(1,25) = 0.9, p = 0.3, ηp^2^ = 0.04; Females: SD F(1,24) = 10.6, p = 0.003, ηp^2^ = 0.31; HFD F(1,24) = 6.4, p = 0.02, ηp^2^ = 0.21; vHFD F(1,24) = 4.5, p = 0.04, ηp^2^ = 0.16). SD, 8M/8F (black/gray respectively); HFD, 11M/10F (dark and light pink); vHFD, 9M/9F (dark and light green). Data are presented as mean ± SEM (bars or closed circles) with individual values (thin lines). *, ** p < 0.05 and 0.01 respectively (Planned comparisons Valued vs Devalued with Bonferroni’s correction). #, ##, ### p < 0.05, 0.01 and 0.001 respectively (one sample t-test versus 0.5). Full statistical reporting is provided in **Supplemental Table 2**. Mouse illustration from NIAID NIH BioArt Source (bioart.niaid.nih.gov/bioart/20).

Finally, we also tested their ability to update action-outcome relationships using an instrumental contingency degradation procedure during which non-expected food rewards were introduced (**Figure 4A**). Despite reversal and degradation procedures both requiring the updating of previous acquired action-outcome relationships, contingency degradation relies on moving from a positive to a null relationship between an action and its former consequences. Both male and female SD mice decreased their levels of response when the instrumental contingency was degraded, compared to mice with intact contingency. HFD-exposed males similarly showed sensitivity to contingency degradation. In contrast, HFD-exposed females maintained a stable response level whether the instrumental contingency being maintained or degraded. Lastly, both male and female vHFD-exposed groups are also insensitive to changes in instrumental contingency, maintaining a high level of response across the sessions (**Figure 4B**). A similar behavioral pattern was observed during the subsequent choice test (**Figure 4C**), clearly demonstrating an impairment in these groups’ ability to correctly update action-outcome relationships.

**Figure 4.**
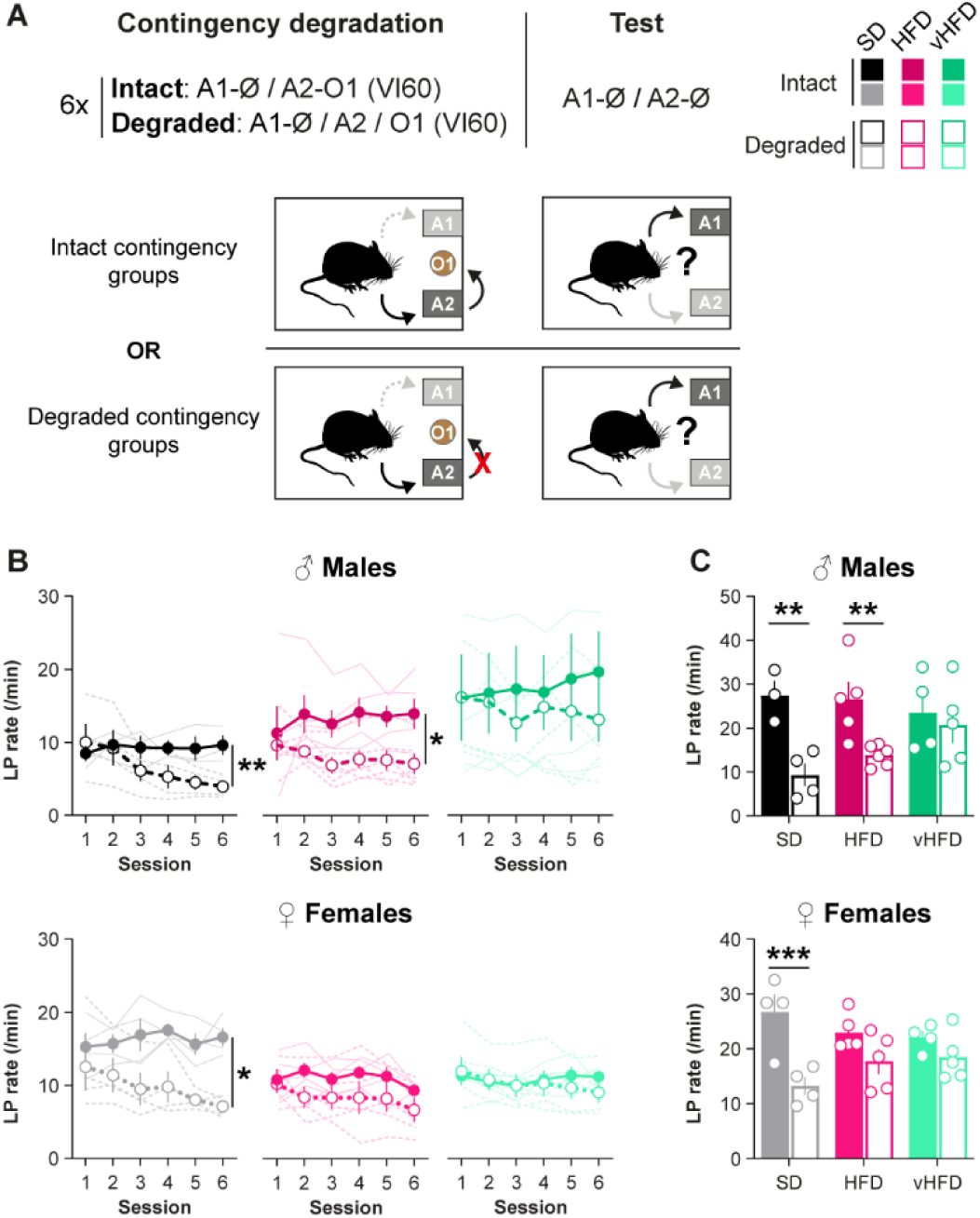
Adolescent obesogenic diets alter adaptation to the degradation of action-outcome contingency at adulthood in a sex-dependent manner. A. Experimental timeline for instrumental retraining followed by contingency degradation and extinction test. For half of the mice, the contingency A2 à O was degraded (A2 / O) for 6 sessions, whereas this contingency remains intact for the rest of the mice. B. Response rate for mice with the non-degraded (intact, solid lines) or degraded (dashed lines) instrumental contingency (two-way ANOVA; Contingency effect; Males: CD F(1, 5) = 28.4, p = 0.003,η_p_^2^ = 0.85; HFD F(1, 9) = 5.4, p = 0.04, η_p_^2^ = 0.38; vHFD F(1, 7) = 0.3, p = 0.6, η_p_^2^ = 0.04 / Females: F(1, 6) = 8.4, p = 0.03, η_p_^2^ = 0.58; HFD F(1, 8) = 2.2, p = 0.2, η_p_^2^ = 0.22; vHFD F(1, 7) = 0.1, p = 0.8 η_p_^2^ = 0.01). C. Response rate on the active lever during the subsequent choice test in extinction (Planned comparisons with Bonferroni’s correction; Males: SD F(1,21) = 11.1, p = 0.003, η_p_^2^ = 0.35; HFD F(1,21) = 8.7, p = 0.008, η_p_^2^ = 0.29; vHFD F(1,21) = 0.3, p = 0.6, η_p_^2^ = 0.02 / Females: SD F (1,21)= 19.2, p < 0.001, η_p_^2^ = 0.48; HFD F(1,21) = 3.8, p = 0.07, η_p_^2^ = 0.15; vHFD F(1,21) = 1.5, p = 0.2, η_p_^2^ = 0.07). SD, 7M/8F (black/gray respectively); HFD, 11M/10F (dark and light pink); vHFD, 9M/9F (dark and light green). Data are presented as mean ± SEM (bars or large circles) with individual values (small open circles or thin lines). *, **, *** p < 0.05, 0.01 and 0.001 respectively Contingency degradation effect (two-way ANOVA followed by Bonferroni’s *post hoc* tests / Planned comparisons Intact vs Degraded with Bonferroni’s correction). Full statistical reporting is provided in Supplemental Table 2. Mouse illustration from NIAID NIH BioArt Source (bioart.niaid.nih.gov/bioart/20).

These results demonstrate that exposure to vHFD during adolescence impairs the ability to detect changes in action-outcome relationships in both sexes. However, exposure to a moderately high-fat diet during adolescence appeared to specifically alter this cognitive process in females, highlighting again sex differences in the impact of diet on cognition.

## DISCUSSION

The overconsumption of highly palatable energy-dense foods is one of the main drivers underlying the current rising levels of obesity, especially in younger populations. Using a diet-induced obesity mouse model restricted to adolescence, we reveal long-lasting alterations of decision-making processes and behavioral flexibility. More importantly, we demonstrate that these effects are dependent on sex and on the nutrient (fat) content of the obesogenic diet, highlighting these as critical factors.

The impact of high-fat diets on body weight and metabolic health is highly dependent on animal sex and dietary fatty acid levels. Consumption of food with high fat content during adolescence increases body weight and fat mass levels and dysregulates glucose homeostasis, consistent with previous studies [39,40]. As expected, mice fed with a diet containing very high levels of fat (60%) had the highest body weight gain associated with the accumulation of fat mass and alterations of metabolic health. The relevance of such diet with human physiology and consumption is debated [40]. However, vHFD rapidly induces obesity and metabolic alteration which represents a massive advantage for developmental approaches such as the one presented here. On the other hand, mice fed with diet containing moderate levels of fatty acids (45%) showed a relatively limited weight gain and proportion of fat mass, even after 5 weeks, an effect mainly observed in males. This is consistent with the sex difference in fat accumulation and obesity development reported in humans and animal models, with males being more vulnerable [37,38].

Our results are in stark contrast with the effects of high fat diets given in adulthood, where even moderate high fat diet produces rapid weight gain and obesity [39,40,47], suggesting that the interaction between food intake and metabolic effects are different in adolescence as we previously observed with protein restriction [48]. Interestingly, these diet effects were not associated with major changes in energy intake. It is also important to note that these body weight and metabolic changes did not persist in adulthood, suggesting that behavioral alterations could not be fully attributed to broader physiological changes usually observed in chronically obese animals (see also **Supplemental Figure 5**). Without access to individual food intake, it remains however difficult to fully dissociate effects from total energy intake and those from specific nutrients. SD, HFD and vHFD have major differences in macronutrients content but also in the nature of these macronutrients, with high-fat diets containing high levels of saturated and monounsaturated fatty acids (lard), sucrose and maltodextrin. Thus, the direct comparison between the effects of the two diets presented here highlights the importance of considering the nutrient content, more than absolute calories, in the regulation of body weight and in the effects of diet exposure on cognition. The precise impact of each macronutrient and their balance remains to be determined, requiring custom-made unbalanced and control diets with matching composition [39,40].

Numerous studies suggest that goal-directed and habitual controlling systems collaborate and interact from early learning stages to control action selection and performance [49]. To investigate how adolescent exposure to obesogenic diets may accelerate habitual control we used an extended training protocol with VI reinforcement schedules. Both overtraining and interval schedules are known to promote habitual behavior by inducing a low correlation between instrumental performance and reward rate [15,16,49]. Here we showed similar instrumental acquisition and performance independently of diet exposure, as reported in previous works [20,23,50]. However, others have shown decreased instrumental responding after diet-induced obesity attributed to a decreased value associated with non-obesogenic diets or differences in learning processes [24,27,51,52,29,53]. This discrepancy might be explained by differences in experimental procedures (animal models, diet-induced obesity models, food rewards).

Using repeated outcome devaluation tests performed at different learning stages [43,44], we observed as expected a progressive transition from goal-directed (sensitive to outcome devaluation) to habitual (insensitive to outcome devaluation) control of actions in control males. Intriguingly, we did not observe such transition in control females, even after extended training (26 days, ∼780 action-outcome pairings). Increasing evidence shows sex-specific strategies in decision-making tasks [30,32,33] but only a limited number of studies directly compared males and females in the study of action control. In contrast to our results, Toufexis and colleagues previously showed that females rats exhibit a faster transition to habitual behavior [54,55], while other studies showed similar impairment of goal-directed control after exposure to palatable food [25]. As for the learning phase, this could result from methodological differences between studies (e.g. animal species, instrumental operandum, outcome devaluation procedures), warranting further investigations. Using two levers (active and inactive) in our task for instance may prevent, or at least slow down, the transition to habitual behavior as previously shown with choice procedures [14]. Thus, we cannot fully exclude the possibility that both females SD and HFD mice, given additional training, would eventually become sensitive to outcome devaluation.

Furthermore, it is important to note that the instrumental performance during the tests progressively decreased in all groups despite similar performance during reacquisition, which might result in a floor effect and mask potential sensitivity to outcome devaluation. This pattern has been observed by others [44] and might result from the increased competition between the original instrumental learning and the inhibitory extinction learning occurring during each tests [56]. Nevertheless, the maintained sensitivity to outcome devaluation observed after extended training in some female groups strongly suggest that such competition does not impact the expression of goal-directed actions when present.

We demonstrate that chronic consumption of HFDs decreases sensitivity to outcome devaluation, suggesting impaired flexible action control, consistent with previous work in humans and animal models [20–25]. We also reveal important sex differences in HFD-induced alterations of action control. Insensitivity to outcome devaluation occurs after a moderate amount of training for both HFD and vHFD-exposed males but appears much earlier in vHFD-exposed female mice. Moreover, the sensitivity of HFD-exposed females to outcome devaluation is in striking contrast with the alteration observed in HFD males. Importantly these effects cannot be explained by differences in task learning, motivation for food reward or in efficiency of sensory-specific satiety, which were similar in all groups. Based on this, our results strongly support that obesogenic diets alter their ability to retrieve outcome representation to control their actions. However, this effect may be dependent on diet-induced changes in body weight and metabolic health, as only HFD-exposed males and all vHFD-exposed mice display these behavioral deficits. Body weight gain at the end of adolescence and fat mass levels were only weakly correlated with behavioral performance during outcome devaluation (**Supplemental Figure 5** and **Supplemental Table 4**). Moreover, all mice exhibited similar body weights by the end of behavioral testing. This suggests that, while we cannot exclude overweight/obesity participating to the long-lasting alterations of action control reported here, it does not fully explain the complex diet- and sex-dependent pattern observed.

The adaptive control of actions also requires the updating and flexible use of action-outcome relationships, a process which is often overlooked in the literature looking at food impact on decision-making. We tested this using instrumental contingency reversal and degradation procedures. Instrumental reversal followed by outcome devaluation requires animals to encode and use a new set of A-O associations. Consistently, all SD mice showed goal-directed control in accordance with the new instrumental contingencies, demonstrating that the reversal procedure is sufficient to restore goal-directed control in overtrained mice. Moreover, these groups also showed some form of goal-directed behavior on the previous action, showing that the initial A-O relationship is not forgotten [57,58]. Similar results were observed in male HFD-exposed mice, demonstrating that this form of behavioral flexibility was intact. The reversal procedure is sufficient to restore goal-directed control even in overtrained mice. In contrast, goal-directed control was impaired in HFD-exposed females and in all vHFD-exposed animal, suggesting complex interactions between diet and sex in different action control processes.

Interestingly, a similar pattern of results was observed in response to contingency degradation. While also involving the updating of A-O relationships as reversal learning, contingency degradation specifically requires the animal to learn that there is now no clear relationship between their action and the reward delivery. This increased uncertainty driven by the delivery of rewards non-contingent with action performance may lead to a decrease in action value and competitions with other forms of associations [14,59–62], which is not the case with reversal procedure. There is a very limited number of studies comparing the two processes and it is likely that they are supported by mostly non-overlapping brain circuits [57,63,64]. Here our results demonstrate that both processes are impacted according to specific diets. To our knowledge, this is the first study showing such diet-induced deficits in females. Interestingly, behavioral adaptation to reversal and contingency degradation was only weakly correlated to body weight parameters (**Supplemental Figure 5** and **Supplemental Table 4**), suggesting once again that alterations of action control processes observed here are not only supported by changes in body weight.

Our experimental design allowed us to compare these multiple processes and the diet impact on the same animals. While this within-subject design has been used in other studies [44,57], repeated testing may represent an important limitation which needs to be considered. Previous work has shown that behavioral test batteries and training history might impact some behavioral responses [65]. Here, mice were tested multiple times including in extinction conditions and with several manipulations of action-outcome relationships (reversal, contingency degradation). Mice were systematically retrained between the different tests to reach similar baseline performance levels, and they showed similar response extinction during tests (*data not shown*). However, it remains to be determined if similar effects would be observed on separated cohorts or with different testing orders (e.g. contingency degradation before response reversal [57]). While our *a priori* comparisons revealed robust sensitivity (or lack of) to outcome devaluation and contingency degradation in our different groups, our results also have to be interpreted with caution as statistical analyses did not reveal any significant interactions in the performance between diets and testing conditions.

It is important to note that, in the present study, animals were tested several weeks after being switched to a healthier diet. Most studies using early life diet exposure maintained subjects under obesogenic diets during behavioral investigation which may induce confounding impact of early exposure and the ongoing development of obesity [28,29,53,66,67]. On the other hand, studies in which animals were switched to a healthy diet report contradictory results associated with either a rescue [68–70] or with long-term deficits [52,71–73] of cognitive processes, depending of the diet intervention. Here we clearly demonstrate long-lasting impact of cognitive processes underlying action control, which may support poor long-term outcomes and frequent relapse to unhealthy dietary habits often observe in obese people [11,12].

The neurobiological bases supporting these different diet-induced behavioral alterations remain to be explored. The flexible control of action is supported by distinct, but interacting, cortico-subcortical circuits determining the balance between goal-directed and habitual control [16–19]. Among these circuits, previous studies have shown that prefrontal [10,24,74] and hippocampal [28,66,67,75] regions are especially sensitive to the effects of palatable foods, even at adulthood. As the prefrontal cortex, the hippocampal formation and their neuromodulatory systems end their development during adolescence [2–7], future studies could explore if obesogenic diets differentially impact them depending on diet nutrient content, related metabolic changes and sex hormones. This might explain the diversity of cognitive impairment we reported here [76].

In summary, we reveal that the chronic consumption of energy-rich obesogenic diets during adolescence leads to long-lasting impairments in multiple cognitive processes underlying the flexible control of decision-making, which may support long-term vulnerability to obesity development and food-related disorders. Importantly, we show that these impairments are sex-dependent and highly influenced by the nutrient content of the diets and their short-term effects on metabolic health. To determine how such changes can be prevented and to develop new therapeutic approaches, future studies will need to dissect precisely how unbalanced dietary habits impact the dynamic remodeling of brain circuits occurring during adolescence.

## DATA AVAILABILITY

Upon publication, all data analyzed in this paper will be available on Figshare https://doi.org/10.6084/m9.figshare.31333456

## ACKNOWLEDGEMENTS

The authors would like to acknowledge the help and support from the staff of the Medical Research Facility, University of Aberdeen for technical support and the care of experimental animals, and Prof. Lora Heisler for sharing equipment used in this study. The authors wish to thank Shauna Parkes, Guillaume Ferreira, James McCutcheon and Etienne Coutureau for helpful discussion and comments on this manuscript.

## AUTHOR CONTRIBUTIONS

DM: Investigation, Writing – Review & Editing; SR: Investigation, Writing – Review & Editing; RTW: Investigation, Validation, Writing – Review & Editing; AR: Investigation, Writing – Review & Editing; KZP: Conceptualization, Formal Analysis, Methodology, Visualization, Writing – Review & Editing; FN: Conceptualization, Data Curation, Formal Analysis, Funding Acquisition, Investigation, Methodology, Project Administration, Supervision, Visualization; Writing – Original Draft Preparation, Writing – Review & Editing.

All authors approved the final version of the manuscript.

## Supplemental Figures and Tables

**Supplemental Figure 1.**
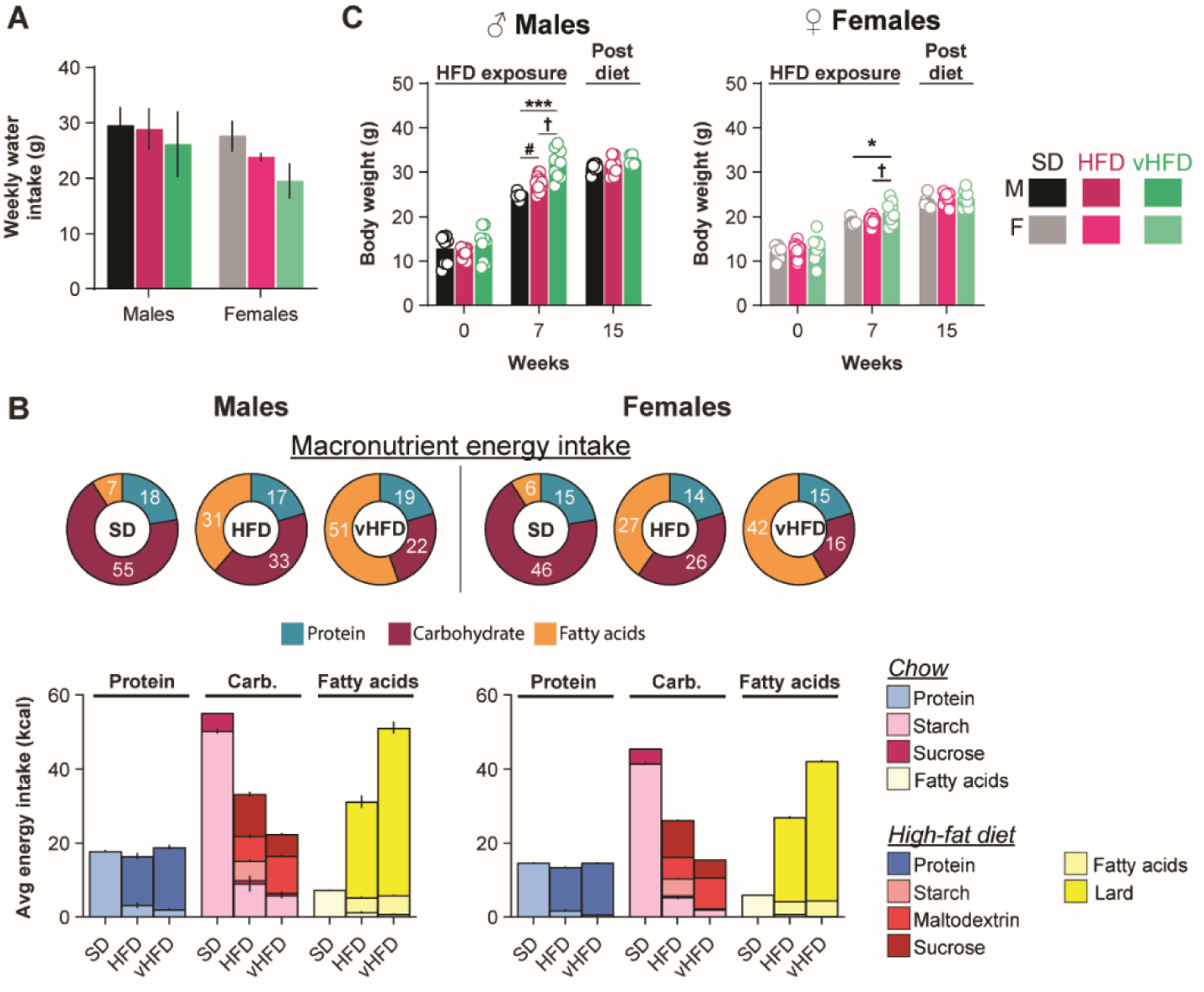
**A.** Weekly water intake (g, estimated from cage consumption) is similar across all groups. **B.** Macronutrient breakdown for average weekly energy intake (in kcal) during diet exposure. Nutrient type is specifically color coded depending on the food source (chow or high-fat diet). **C.** Body weight at the start (0; Males: F_(2, 25)_ = 0.9, p = 0.4, η^2^ = 0.07 / Females: F_(2, 26)_ = 0.4, p = 0.7, η^2^ = 0.03), the end of diet exposure (5 weeks; Males: F_(2,25)_ = 22.6, p < 0.001, η^2^ = 0.64 / Females: F_(2, 26)_ = 5.3, p = 0.01, η^2^ = 0.29) and at the end of behavioral testing (15 weeks; Males: F_(2, 22)_ = 1.5, p = 0.3, η^2^ = 0.12 / Females: F^(2, 24)^ = 1.3, p = 0.3, η^2^ = 0.10). SD, 8 M/8F (black/gray respectively); HFD, 11M/12F (dark and light pink); vHFD, 9M/9F (dark and light green). Data are presented as mean ± SEM with individual values (open circles). * p < 0.05, *** p < 0.001 SD vs vHFD; # p < 0.05 SD vs HFD; † p < 0.05 HFD vs vHFD (one-way or two-way ANOVA followed by Bonferroni’s *post hoc* tests). Full statistical reporting is provided in **Supplemental Table 3**.

**Supplemental Figure 2.**
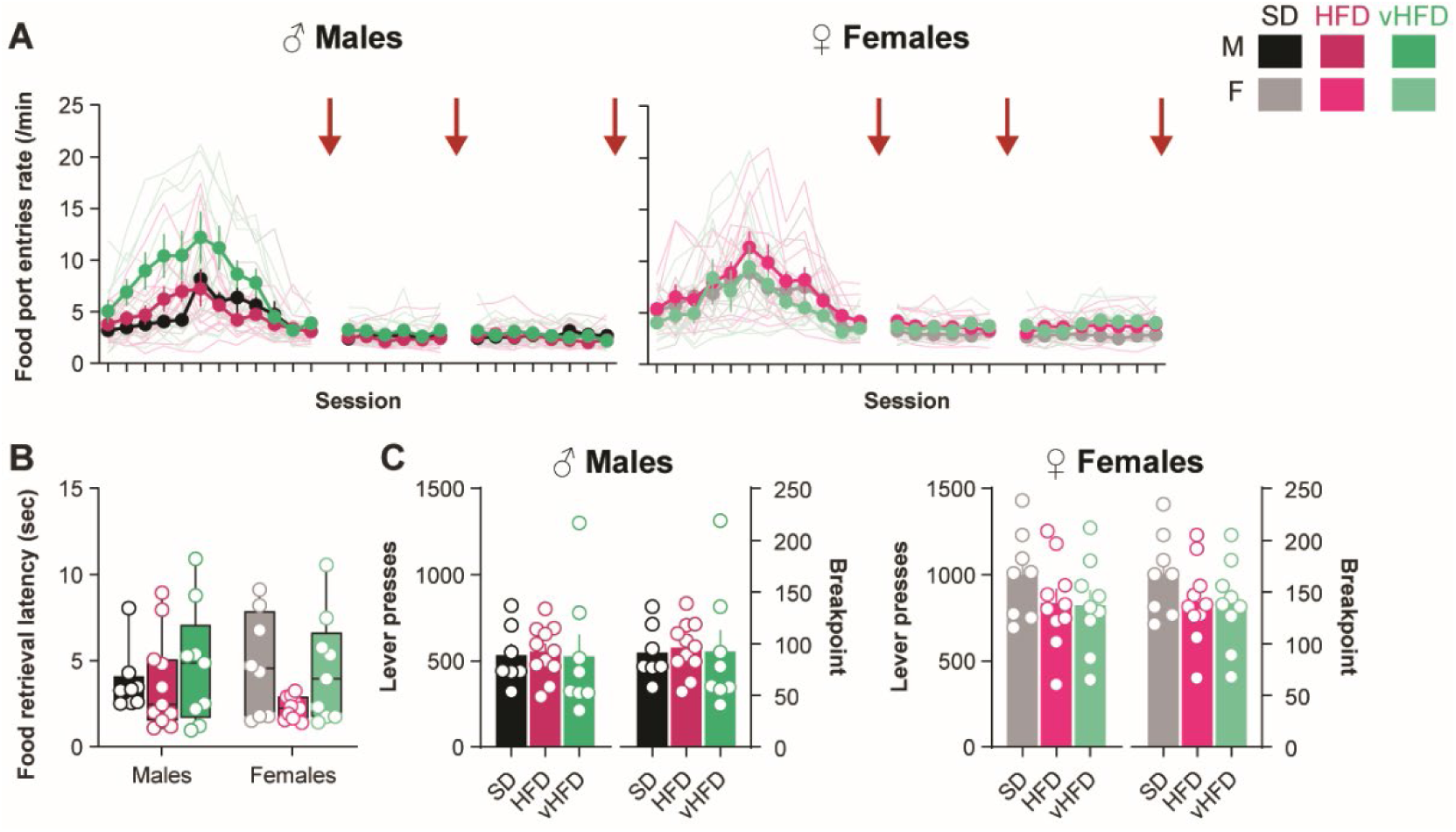
**A.** Food port entries response rate across initial training. Red arrows indicate the timing of the outcome devaluation tests. **B.** Average food retrieval latency during initial instrumental training. **C.** Progressive ratio task. Average total lever presses and breakpoint for male and female groups. SD, 7-9M/8-9F (black/gray respectively); HFD, 11M/10-11F (dark and light pink); vHFD, 8-9M/9F (dark and light green). Data are presented as mean ± SEM (bars or closed circles) with individual values (open circles or thin lines). Full statistical reporting is provided in **Supplemental Table 3**.

**Supplemental Figure 3.**
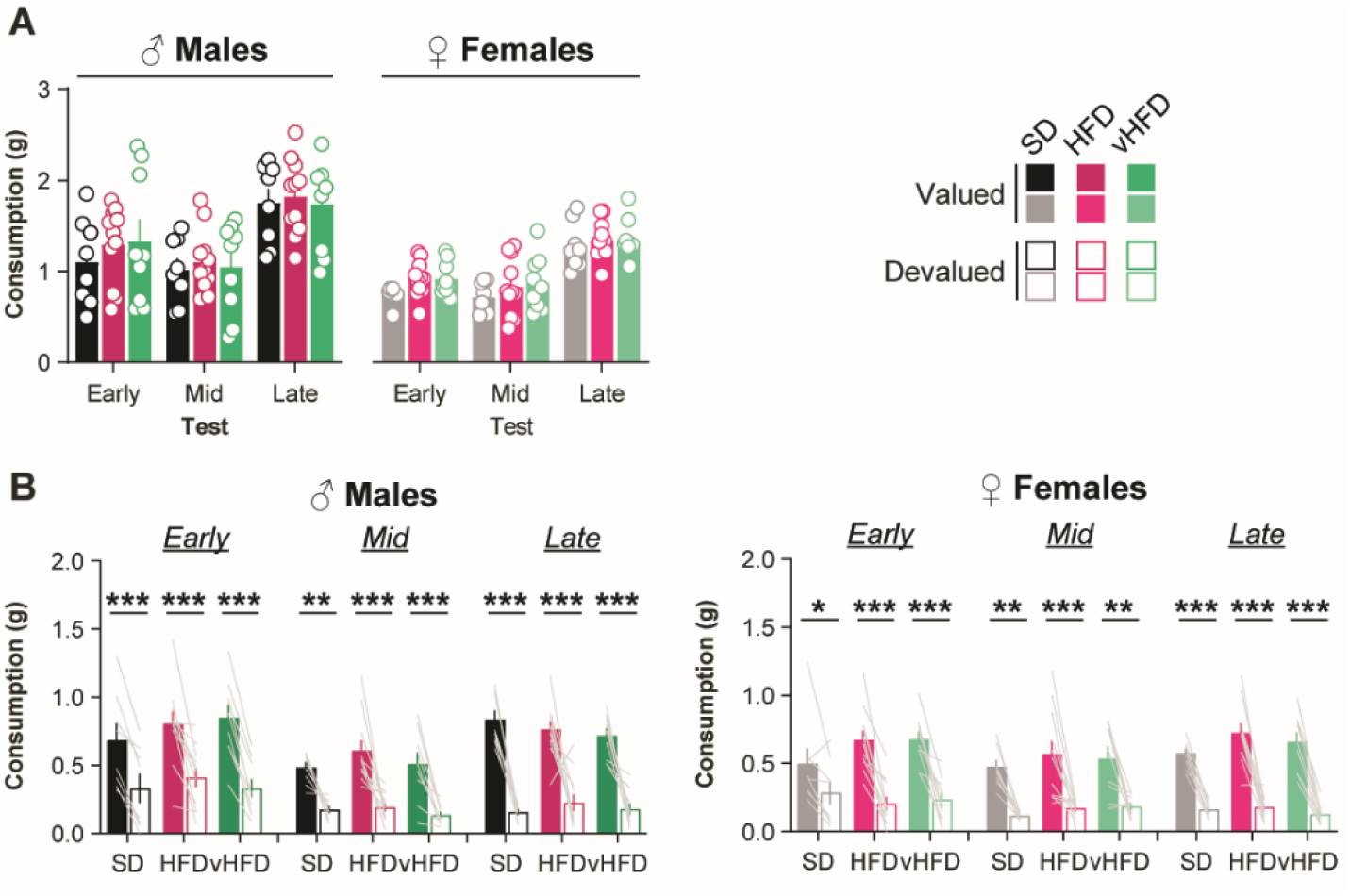
**A.** Food consumption (g) during the satiety phase of early, mid and late outcome devaluation tests. **B.** Food consumption (g) the choice consumption tests after each outcome devaluation test. SD, 8M/8F (black/gray respectively); HFD, 11M/11F (dark and light pink); vHFD, 9M/9F (dark and light green). Data are presented as mean ± SEM (bars or closed circles) with individual values (open circles or thin lines). *, **, *** p < 0.05, 0.01 and 0.001 respectively Devaluation effect (two-way ANOVA followed by Bonferroni’s *post hoc* tests). Full statistical reporting is provided in **Supplemental Table 3**.

**Supplemental Figure 4.**
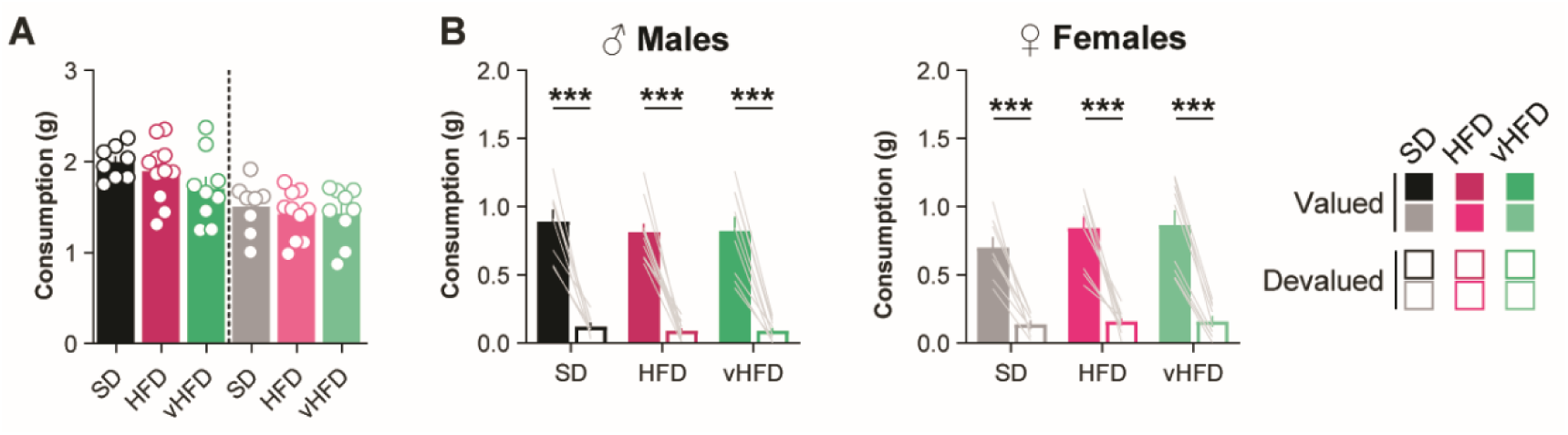
**A.** Food consumption (g) during the outcome devaluation tests after reversal learning. **B.** Food consumption (g) the choice consumption tests after the outcome devaluation test. SD, 8M/8F (black/gray respectively); HFD, 11M/10F (dark and light pink); vHFD, 9M/9F (dark and light green). Data are presented as mean ± SEM (bars or closed circles) with individual values (open circles or thin lines). *, **, *** p < 0.05, 0.01 and 0.001 respectively Devaluation effect (two-way ANOVA followed by Bonferroni’s *post hoc* tests). Full statistical reporting is provided in **Supplemental Table 3**.

**Supplemental Figure 5.**
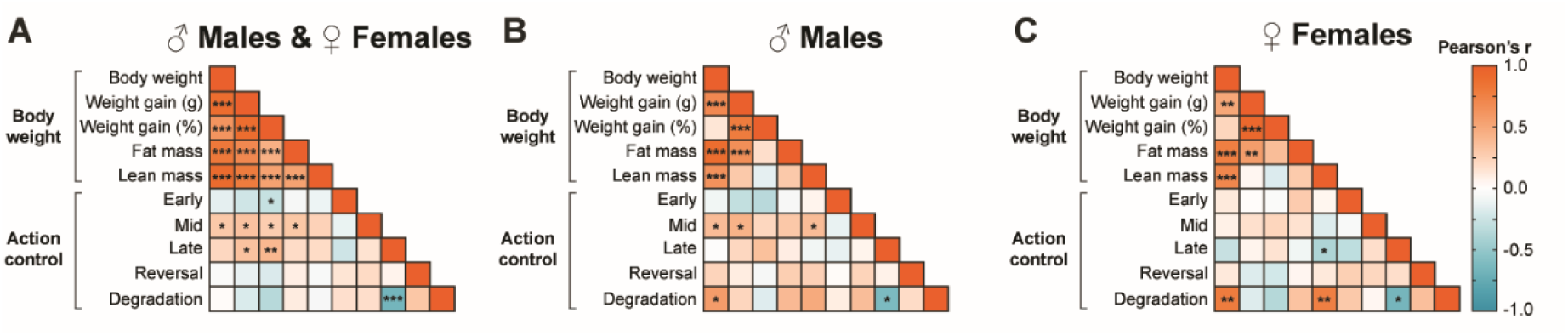
Correlation matrices between body weight measures and action control performance. Heatmaps showing the relationship (Pearson’s r) between body weight at the end of diet exposure, weight gain (g), weight gain (% starting body weight), fat and lean mass (g), Outcome Devaluation indexes (Early, Mid, Late, Reversal; Response rate Devalued/Valued+Devalued) and Contingency Degradation index (Response rate Degraded/Degraded+Baseline) for all mice (A), Males (B) and Females (C). *, **, *** p < 0.05, 0.01 and 0.001 respectively. Full statistical reporting is provided in **Supplemental Table 4**.

**Supplemental Table 1.** Macronutrient breakdown of SD, HFD and vHFD.

|  | SD<br>(CMR,<br>SDS) | HFD<br>(D12451,<br>Research<br>Diets) | vHFD<br>(D12492,<br>Research<br>Diets) |
| --- | --- | --- | --- |
|  | 3.6<br>kcal/g | 4.73<br>kcal/g | 5.24<br>kcal/g |
| **<u>Protein</u>** | **22** | **20** | **20** |
| **<u>Carbohydrate</u>** | **69** | **35** | **20** |
| *Starch &*<br>*fibres* | 65 | 7 | 0 |
| *Sucrose* | 4 | 10 | 7 |
| *Maltodextrin* | 0 | 18 | 13 |
| **<u>Fat</u>** | **9** | **45** | **60** |
| *Lard* | 54 | 39 | 54 |
| *Other* | 6 | 6 | 6 |

**Supplemental Table 2:**
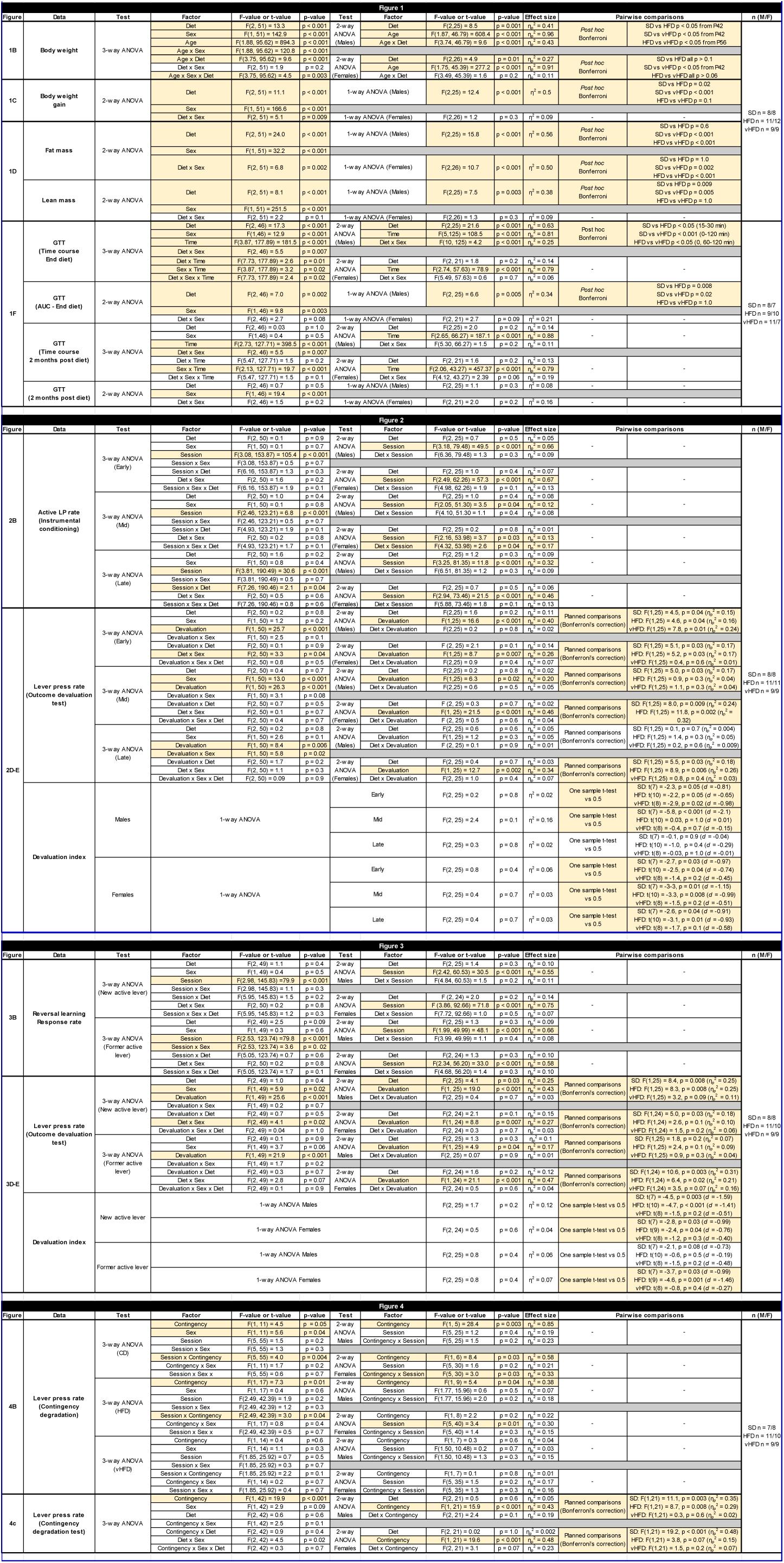
Full statistical report for Figures 1-4.

**Supplemental Table 3:**
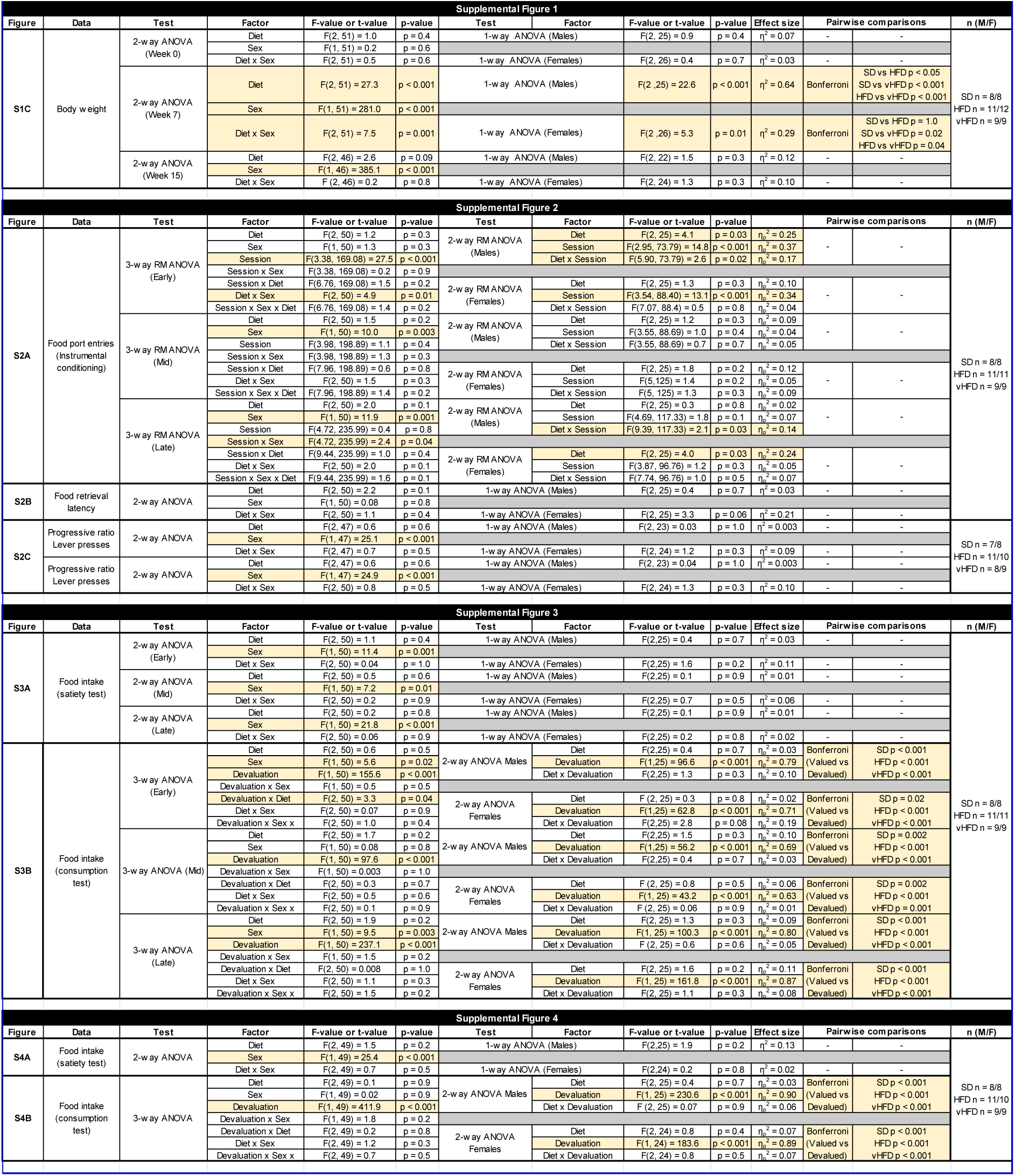
Full statistical report for Supplemental Figures 1-4.

**Supplemental Table 4.** Statistical report for correlation matrices between body weight measures and action control performance (Pearson’s r). *, **, *** p < 0.05, 0.01 and 0.001 respectively.

| A- All mice |  | Test Variables |  |  |  |  |  |  |  |  |  |
| --- | --- | --- | --- | --- | --- | --- | --- | --- | --- | --- | --- |
|  | Test variables | 1 | 2 | 3 | 4 | 5 | 6 | 7 | 8 | 9 | 10 |
| Body weight | 1- Body weight | - |  |  |  |  |  |  |  |  |  |
|  | 2- Weigh gain (g) | 0.893*** | - |  |  |  |  |  |  |  |  |
|  | 3- Weight gain (% baseline) | 0.624*** | 0.898*** | - |  |  |  |  |  |  |  |
|  | 4- Fat mass | 0.815*** | 0.704*** | 0.452*** | - |  |  |  |  |  |  |
|  | 5- Lean mass | 0.908*** | 0.79*** | 0.539*** | 0.540*** | - |  |  |  |  |  |
| Action control | 6- Devaluation - Early | -0.145 | -0.239 | -0.282* | -0.056 | -0.106 | - |  |  |  |  |
|  | 7- Devaluation - Mid | 0.305* | 0.340* | 0.277* | 0.314* | 0.246 | -0.119 | - |  |  |  |
|  | 8- Devaluation - Late | 0.216 | 0.343* | 0.405** | 0.191 | 0.192 | -0.242 | 0.157 | - |  |  |
|  | 9- Devaluation - Reversal | -0.031 | -0.129 | -0.186 | 0.095 | -0.048 | 0.144 | 0.199 | 0.071 | - |  |
|  | 10- Degradation | 0.014 | -0.189 | -0.351 | 0.109 | -0.051 | 0.172 | 0.18 | -0.67*** | 0.315 | - |
| B- Males |  | Test Variables |  |  |  |  |  |  |  |  |  |
|  | Test variables | 1 | 2 | 3 | 4 | 5 | 6 | 7 | 8 | 9 | 10 |
| Body weight | 1- Body weight | - |  |  |  |  |  |  |  |  |  |
|  | 2- Weigh gain (g) | 0.701*** | - |  |  |  |  |  |  |  |  |
|  | 3- Weight gain (% baseline) | 0.138 | 0.787*** | - |  |  |  |  |  |  |  |
|  | 4- Fat mass | 0.892*** | 0.655*** | 0.177 | - |  |  |  |  |  |  |
|  | 5- Lean mass | 0.638*** | 0.298 | -0.142 | 0.300 | - |  |  |  |  |  |
| Action control | 6- Devaluation - Early | -0.088 | -0.303 | -0.344 | -0.053 | 0.063 | - |  |  |  |  |
|  | 7- Devaluation - Mid | 0.38* | 0.438* | 0.221 | 0.31 | 0.379* | -0.124 | - |  |  |  |
|  | 8- Devaluation - Late | -0.015 | 0.219 | 0.357 | 0.102 | -0.081 | -0.133 | 0.067 | - |  |  |
|  | 9- Devaluation - Reversal | 0.236 | 0.106 | -0.056 | 0.269 | 0.258 | 0.018 | 0.242 | 0.081 | - |  |
|  | 10- Degradation | 0.575* | 0.225 | -0.164 | 0.391 | 0.486 | 0.101 | 0.372 | -0.623* | 0.191 | - |
| C- Females |  | Test Variables |  |  |  |  |  |  |  |  |  |
|  | Test variables | 1 | 2 | 3 | 4 | 5 | 6 | 7 | 8 | 9 | 10 |
| Body weight | 1- Body weight | - |  |  |  |  |  |  |  |  |  |
|  | 2- Weigh gain (g) | 0.505** | - |  |  |  |  |  |  |  |  |
|  | 3- Weight gain (% baseline) | 0.231 | 0.937*** | - |  |  |  |  |  |  |  |
|  | 4- Fat mass | 0.770*** | 0.566** | 0.374 | - |  |  |  |  |  |  |
|  | 5- Lean mass | 0.717*** | 0.053 | -0.181 | 0.293 | - |  |  |  |  |  |
| Action control | 6- Devaluation - Early | 0.119 | 0.007 | -0.030 | 0.290 | 0.061 | - |  |  |  |  |
|  | 7- Devaluation - Mid | 0.087 | 0.167 | 0.229 | 0.158 | -0.175 | -0.044 | - |  |  |  |
|  | 8- Devaluation - Late | -0.277 | 0.063 | 0.152 | -0.087 | -0.392* | -0.315 | 0.178 | - |  |  |
|  | 9- Devaluation - Reversal | 0.123 | -0.175 | -0.218 | 0.058 | 0.125 | 0.259 | 0.222 | 0.18 | - |  |
|  | 10- Degradation | 0.787** | -0.174 | -0.362 | 0.339 | 0.762** | 0.299 | 0.048 | -0.659* | 0.360 | - |

